# Summarizing internal dynamics boosts differential analysis and functional interpretation of super enhancers

**DOI:** 10.1101/2021.09.25.461810

**Authors:** Xiang Liu, Bo Zhao, Timothy I. Shaw, Brooke L. Fridley, Derek R. Duckett, Aik-Choon Tan, Mingxiang Teng

## Abstract

Super enhancers (SEs) are broad enhancer domains usually containing multiple constituent enhancers that hold elevated activities in gene regulation. Disruption in one or more constituent enhancers causes aberrant SE activities that lead to gene dysregulation in diseases. To quantify SE aberrations, differential analysis is performed to compare SE activities between cell conditions. The state-of-art strategy in estimating differential SEs relies on overall activities and neglect the changes in length and structure of SEs. Here, we propose a novel computational method to identify differential SEs by weighting the combinatorial effects of constituent-enhancer activities and locations (i.e., internal dynamics). In addition to overall activity changes, our method identified four novel classes of differential SEs with distinct enhancer structural alterations. We demonstrate that these structure alterations hold distinct regulatory impact, such as regulating different number of genes and modulating gene expression with different strengths, highlighting the differentiated regulatory roles of these unexplored SE features. When compared to the existing method, our method showed improved identification of differential SEs that were linked to better discernment of cell-type-specific SE activity and functional interpretation. We implement an R package, *DASE*, to facilitate the use of our method.

## INTRODUCTION

Super enhancers (SEs) were proposed as broad regulatory domains on genome, usually spanning a minimum of thousands of base pairs and consisting of multiple constituent enhancers (CEs) (1). The CEs work together as a unit, instead of separately, to facilitate high enhancer activity, observed as dense enrichment of cell master regulators, coactivators, mediators and chromatin factors at SEs (2). These characteristics were further demonstrated by the fact that, distinct from regular enhancers, SE is specifically linked to gene regulation associated with cell identity and disease mechanisms (3,4).

Recent studies further showed that, beyond the elevated activity, the internal mechanics of SEs also paly critical roles in defining their prominent roles in gene regulation, known as multi-promoter targeting and long-range interactions (5-8). Some SEs form a clear hierarchical structure where hub CEs are responsible for the functional and structural organization of the whole SEs (6,9). Other SEs, in contrary, receive relative balanced contribution from the CEs. In addition, CEs could establish an open chromatin interaction network within individual SEs (7), indicating the internal crosstalk across CEs in orchestrating SEs’ unique functions.

The activity and relations of individual CEs were well appreciated during computational identification of SEs. Existing algorithms usually contain two processing steps (2,10). First, the activity and locations of genome-wide enhancers are inferred through peak detection using chromatin immunoprecipitation sequencing (ChIP-seq) data (11), particularly that measuring the binding of mediators, master regulators, or active histone mark H3K27Ac. Second, the inferred activity and locations are summarized linearly to prioritize broad enhancer regions (2,3), i.e., SEs, that contain densely enriched enhancers with high activities, i.e., the CEs.

However, the organization of CEs were not considered by current approaches in differential analysis of SEs, a key aspect of research interest when comparing across biological conditions (12-16). The alteration of SEs has been found to be highly associated with disease dysregulation and could be used for drug target identification (14-16). These approaches, in which SEs are treated as individual entities, usually identify differential SEs based on binary strategy, which compares the presence and absence of SEs between biological conditions, neglecting the constituent enhancer statuses. Consequently, differential SEs are generated largely depending on the parameters that algorithms utilized to detect super enhancers (2,3). In addition, the sensitivity to detect changes in the local enhancer organization are downplayed within the broad genomic regions occupied by SEs.

Here, we propose a novel computational method to identify differential SEs by summarizing the combinatorial effects of constituent-enhancer activities and locations. In addition to overall activity changes, our method detects four extra differential categories specifically pointing to the internal structural alterations of SEs. We demonstrate the unique characteristics of these differential SE categories using public datasets by linking their altered activity to TF binding and gene expression with 3D chromatin interactions (17-19). The results indicate that each SE category regulates distinct sets of gene targets and their expression. Further, we show that our method maximizes the discernment of cell identities when comparing SE profiles of cell lines from the same cancer type. Our method provides sensitive and biologically meaningful identification of differential SEs, which complements existing understanding of SE dynamics. We implemented an R package, *DASE* (Differential Analysis of Super Enhancers: https://github.com/tenglab/DASE), to facilitate the use of our method.

## RESULTS

### Internal dynamics underlie genome-wide SE differences

SEs usually contain multiple constituent enhancers (CEs) located in close genomic proximity along the genome (2,20). It is important to understand the roles of CEs in contributing to the SE lineage-specificity. We explored the SE profiles for six cancer cell types (**Figure 1** and ***Methods***) using ChIP-seq data of H3K27Ac histone modification from ENCODE project (21). We found that CEs within the same SEs could alter differently across cell types. For example, the CEs, located at two previous reported SE loci responsible for MYC regulation in multiple cancers (22-25), showed divergent activity patterns across the six cell types (**Figure 1a**). We term such divergent alterations between CEs as the internal dynamics of SEs, which underline the individual CE effects on determining the cell-type-specific SE activity.

**Figure 1.**
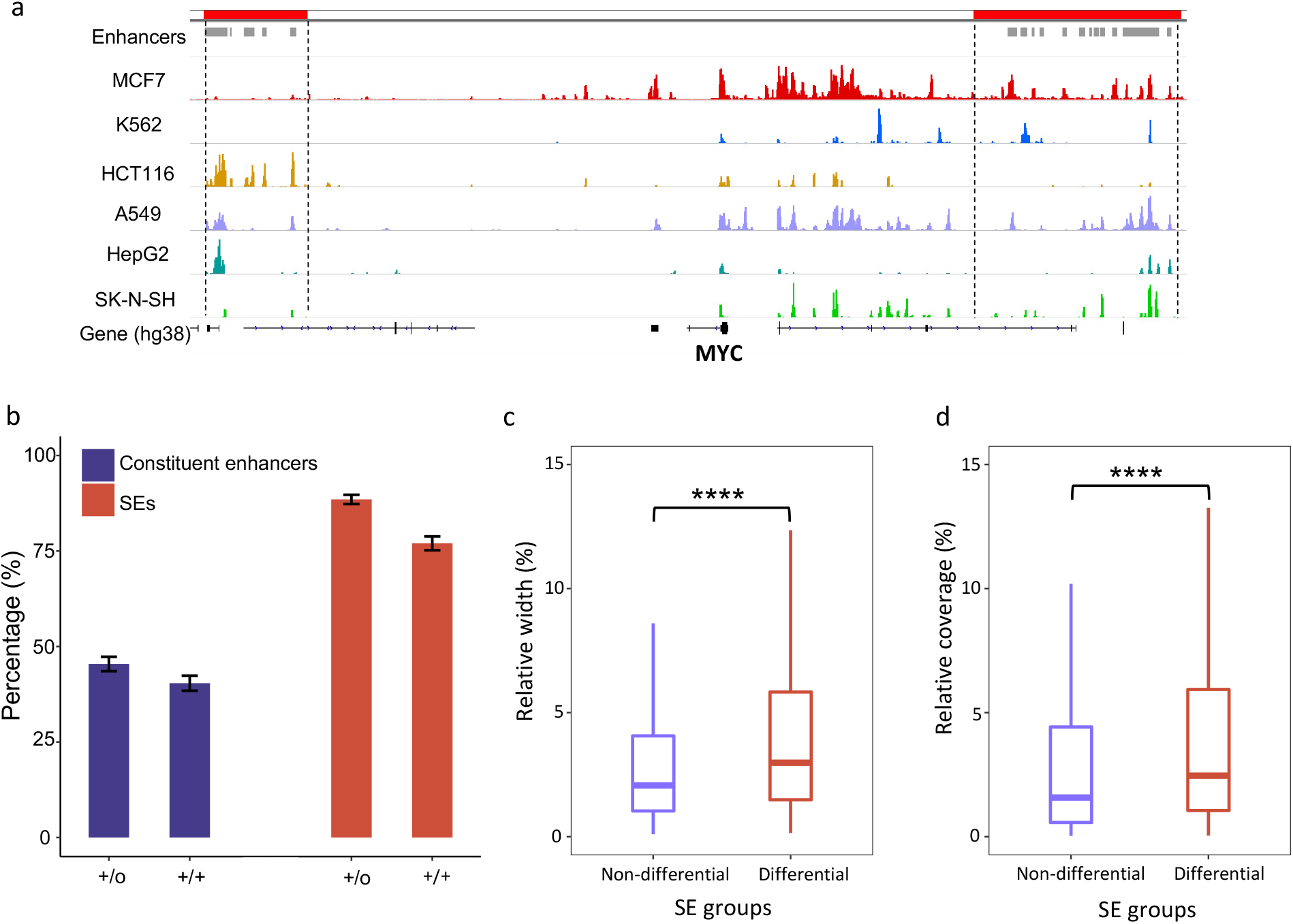
Internal dynamics of super enhancers. **a)**. CEs show frequent alterations across cancer cell types at two reported SE loci associated to MYC regulation. ChIP-seq coverage of H3K27Ac are shown. SEs and CEs are labeled at the top as red and grey bars, separately. **b)**. Frequencies of differential enhancers (fold-change > 4 & q-value < 0.05) and differential SEs (based on binary strategy). Labels on the x-axis indicate features filtered by differential (“+”) or no filtering (“o”), while left and right symbols correspond to filters on CEs and SEs, respectively. **c)**. Relative width of differential CEs from non-differential and differential SEs. Relative width is defined as the percentage of CE width over the summed width from all CEs within a SE. **d)**. Relative coverage of differential CEs from non-differential and differential SEs. Relative coverage is defined as the percentage of CE coverage over the summed coverage from all CEs within a SE.

We then performed pair-wise comparisons of SE profiles across the six cancer cell types. On average, over 40% of CEs showed significant differential activity (fold-change > 4 and q-value < 0.05) that accounts for over 80% of total SEs in these cell types (**Figure 1b**). This indicates that SEs undergo frequent internal alterations across cell types. We further estimated how the alteration of CEs contributed to the overall differences of SEs. Here, we identified differential SEs between cancer cell types using the presence/absence (binary) strategy. A small portion of SEs (∼10%) with significantly altered CEs didn’t show overall differences, while the majority of SEs changed in the same directions as their altered CEs (**Figure 1b**). This implies the divergent influences of CEs on the overall differential statuses of SEs genome-widely.

Next, we examined the characteristics of CEs that might affect their contribution to the overall SE differences. Not surprisingly, the spanning width and regulatory activity of CEs, indicated by H3K27Ac ChIP-seq coverage, showed significant associations with overall SE differences (**Figure 1c-d**). In brief, differential CEs with smaller width or lower activity presented less impacts on the overall statuses of their corresponding SEs. Therefore, we built a model to summarize SE internal dynamics by accounting for these characteristics.

### Modeling internal dynamics leads to distinct patterns of differential SEs

We developed a weighted spline model, implemented as an R package *DASE*, to summarize the internal dynamics into the overall differential statuses of SEs (**Figure 2** and ***Methods***). In brief, differential CEs were first evaluated using existing strategies on detecting differential ChIP-seq peaks (26). Then, a spline was fitted, stratified by enhancer positions, to smooth the differential signals for consecutive CEs. In the smoothing, the activities and width of CEs were taken as fitting weights. Finally, the spline curves were evaluated with permutations to determine reliable differential sub-regions within the SEs, which were further summarized towards the overall differential statuses of SEs.

**Figure 2.**
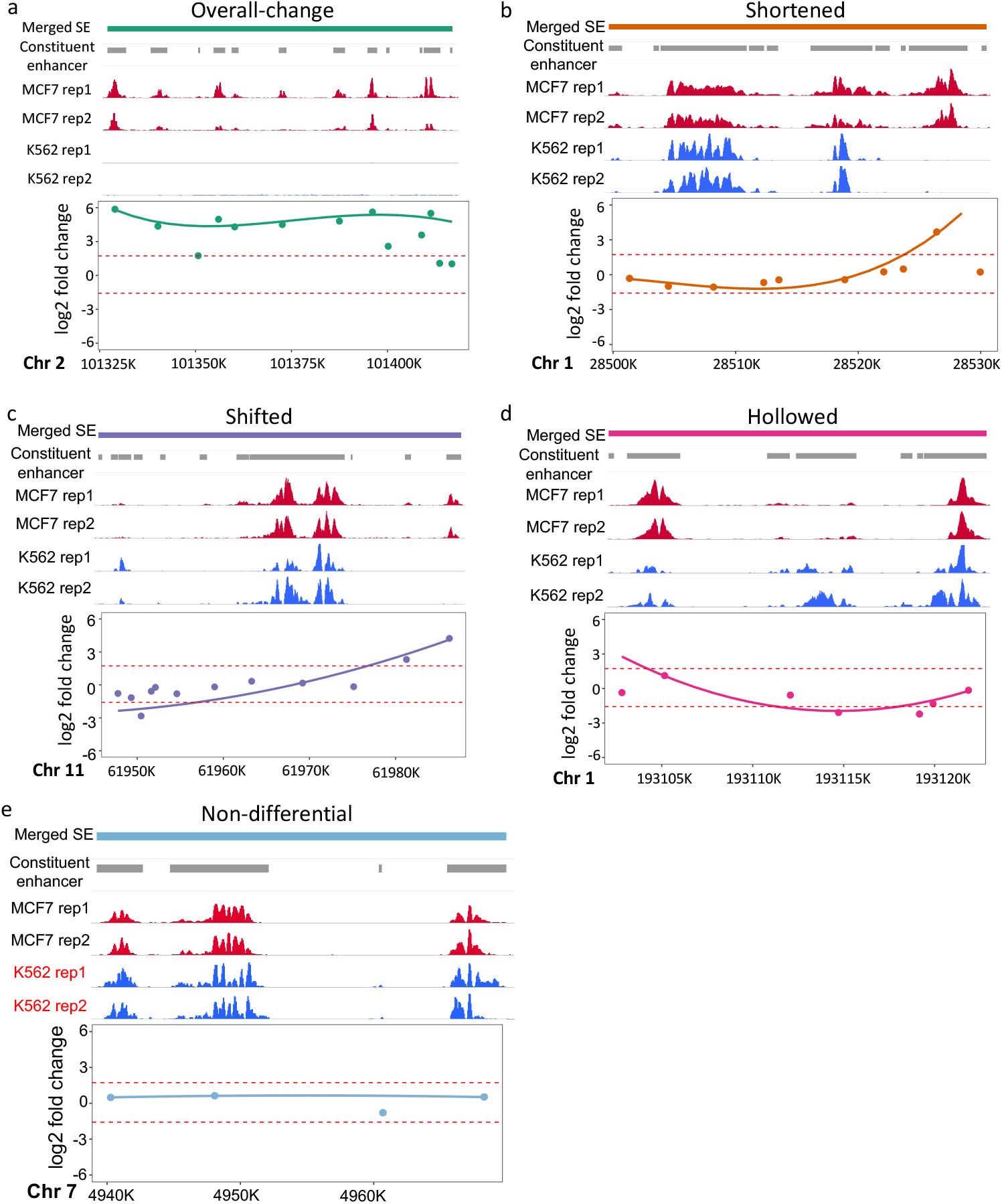
Differential SEs modeled with *DASE*. SE examples are listed with overall-change (**a**), shortened (**b**), shifted (**c**), hollowed (**d**) and non-differential(**e**). In each sub-figure, the upper panel lists in order the SE regions, CEs, H3K27Ac ChIP-seq coverage in two cell types with two replicates. The lower panel shows the fitted b-splines in addition to the original log2 fold-change values for CEs (points). Dashed lines indicate the estimated thresholds from permutation to define differential segments within SE regions. In (**e**), red text indicates the cell line where SE was detected at this locus by the binary strategy.

To illustrate the utility of *DASE*, we compared SE profiles between two cancer cell lines, K562 and MCF7. These two cell lines have high-quality annotation datasets on ENCODE data portal, including transcription factor binding, 3D chromatin interactions and gene expression, to help evaluate the identified differential SEs. *DASE* detected an *overall-change* as well as four novel patterns of differential SEs highlighting the structural alterations within SEs, denoted as *shortened, shifted, hollowed* and *other complex scenarios*, separately (**Figure 2**). *Overall-change* SEs represent significant overall activity alterations (as captured by the binary strategy) as well as consistent altering behavior among CEs. (**Figure 2a**). *Shortened* SEs have significant changes in their sizes by gaining or dismissing CEs on one or both ends of the SEs (**Figure 2b**). *Shifted* SEs have migrated genomic locations without significant size changes (i.e., CEs gained on one end of the SEs and dismissed on the other end) (**Figure 2c**). *Hollowed* SEs represent those with altered CEs in the middle while the two ends remain intact (**Figure 2d**). *Other complex scenario* SEs represent all other complicated or rare cases (**Supplementary Figure 1**). Examples of SE structural alterations reveal that structural differences do not necessarily accompany overall activity differences (**Figure 2a-d**). Together, they provide novel insights to understand SE dynamics between cell conditions. In addition, we note that marginal activity differences that were over-claimed as differential SEs by the binary strategy could be properly corrected by *DASE* (**Figure 2e**).

In total, about 52% of the differential SEs showed *overall-change* between K562 and MCF7, reflecting the distinct chromatin structure underlying each cancer type (**Supplementary Figure 2**). *Shortened* SEs dominated among all types of structural differences (65%), indicating the wide spreading of SE size changes. The other types of structural differences, although not prevailing, represent the diverse dynamics of SE profiles responsible for cell-type-specific gene regulation. We show that those structural differences consistently present in comparisons across more cancer cell types in later sections.

### Diverse differential SEs synergistically build up gene regulation

We further characterized the functional roles of the differential SE patterns in gene regulation. SEs are usually enriched with various transcriptional regulators and cofactors (8), which play critical roles in supporting SE interactions with gene targets (**Figure 3a**). We examined the protein binding profiles across the differential SE patterns. In total, we analyzed 78 transcription factors that have ChIP-seq data available for K562 and MCF7 cell lines by ENCODE project. Transcription factors showed a high correlation with CE activities (**Figure 3b** and **Supplementary Figure 3**), regardless of the differential patterns of the corresponding SEs (**Supplementary Figure 4**), suggesting different patterns of SE alterations share similar mechanisms in recruiting transcription factors. Among these transcription factors, two clear modes of enrichment were identified (**Figure 3b** and **Supplementary Figure 3**): 1) those enriched at active CEs in both K562 and MCF7 cell lines (e.g., MBD2), suggesting that these transcription factors are involved in maintaining key cell functions; 2) those enriched at active CEs in only one cell type but not the other (e.g., NFRKB and FOXA1), indicating their roles in cell-type-specific gene regulation. This suggests different types of SE alterations are involved in both cell-type-specific and housekeeping-related regulation.

**Figure 3.**
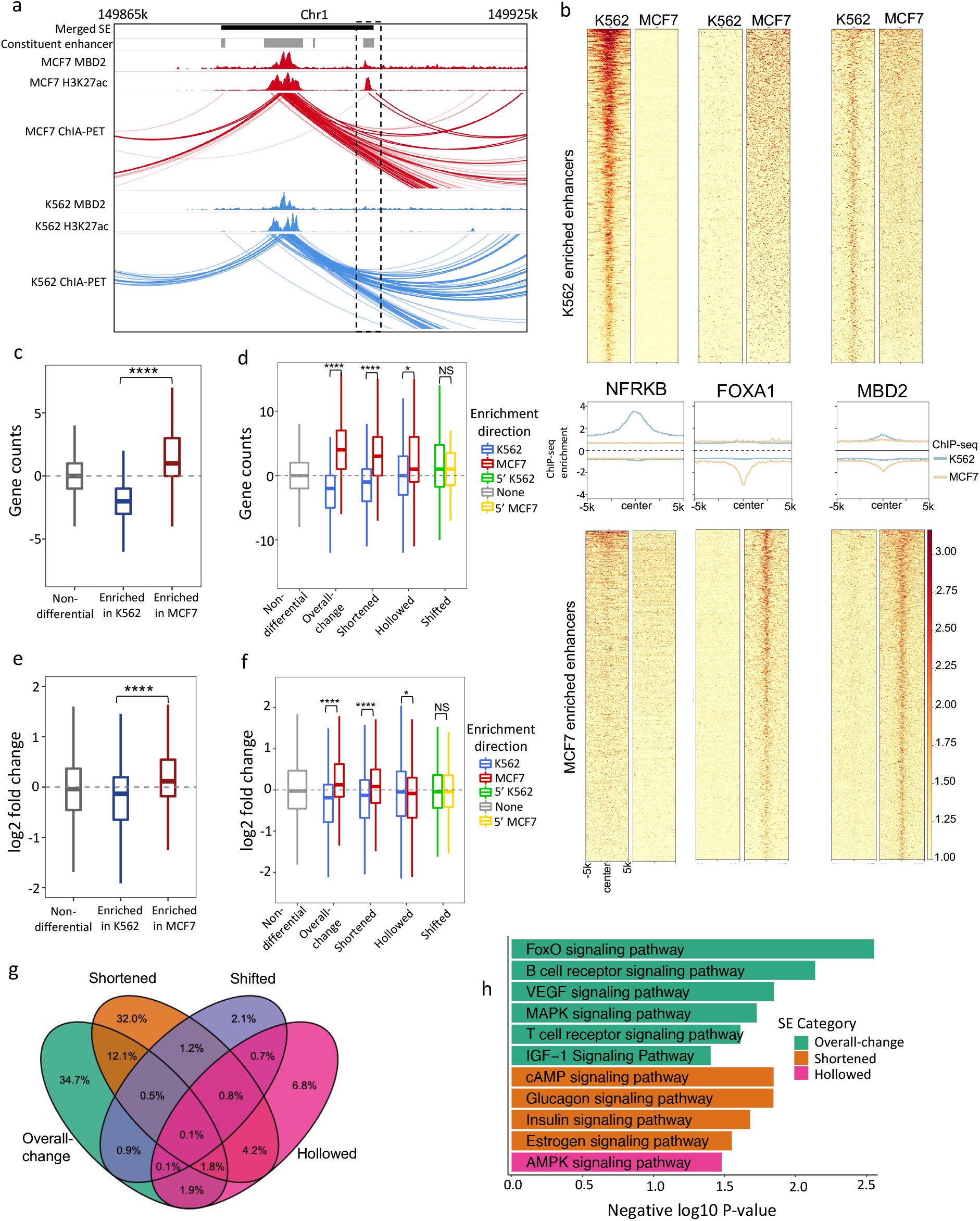
Differential SE categories linked to distinct regulatory features. **a)**. An example shows transcription factor in aiding SE-gene regulation. **b)**. Differential CEs enriched with cell-type-specific (NKRKB & FOXA1) and key-function (MBD2) related transcription factors. Top panels are transcription factor signals at CEs active in K562. Bottom panels are transcription factor signals at CEs enriched in MCF7. Middle panels are the aggregated binding signals from the top and bottom panels with blue and orange lines indicating signals in K562 and MCF7, separately. The signs of y-axises in the middle panels represent enrichment directions. **c)**. Differences on the linked gene numbers between MCF7 and K562 by differential CEs, which are grouped by their altering statuses. Gene targets are identified based on chromatin contacts from POLR2A ChIA-PET data. **d)**. Differences on the linked gene numbers between MCF7 and K562 by differential SEs, which are grouped by their differential patterns. For each differential SE category, SEs are separated into two sub-groups based on their enrichment directions. **e)**. log2 fold-change of the linked genes between MCF7 and K562 by differential CEs. **f**. log2 fold-change of the linked genes between MCF7 and K562 by differential SEs. For each differential SE category, genes are separated into two groups by the enrichment directions of their linked SEs. **g**. Overlaps of the linked genes by four differential SE categories between MCF7 and K562. **h**. KEGG signaling pathways (p < 0.05) uniquely associated to each differential SE category.

**Figure 4.**
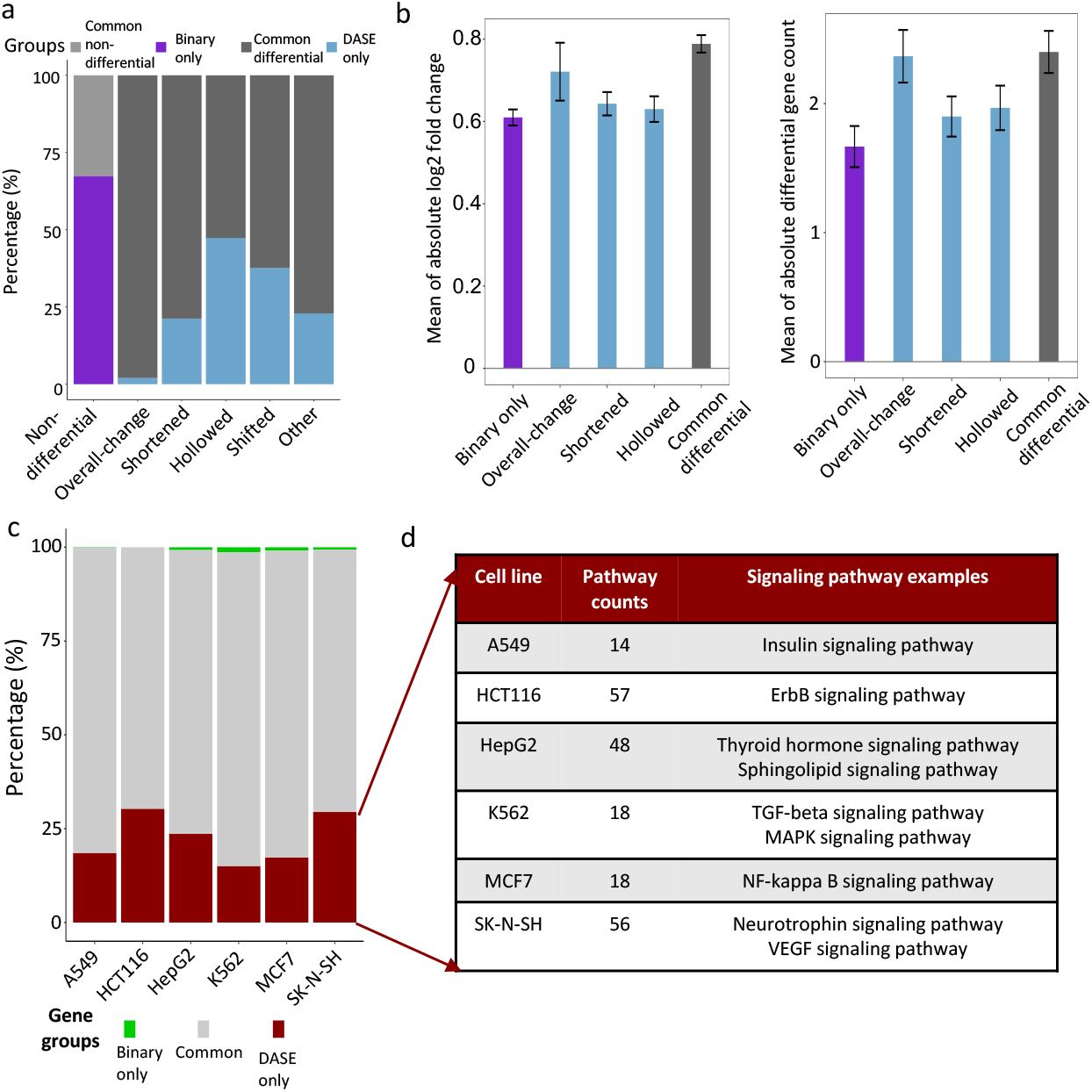
Comparisons between *DASE* and the binary strategy. **a)**. Average discrepancies between *DASE* and the binary strategy across pair-wise comparisons of the six cancer cell types. Light grey: common non-differentials by *DASE* and the binary strategy; Dark grey: common differentials; purple: differential only by the binary strategy; blue: differential only by *DASE*. X-axis is labels with SE categories based on *DASE*. **b)**. Impact on gene expression and linked gene numbers by the discrepant differential SEs between *DASE* and the binary strategy. Bars indicate the mean of log2 fold-change of gene expression or changes of linked gene numbers in each pair-wise comparison and error bars represent the standard error of the means. Results from the two altering directions of the gene effects are merged based on absolute values. **c)**. Genes uniquely linked to SEs with increased activity (overall increased, lengthened or hollowed with increased CEs) in one cancer cell type compared to the other five cells. Green: genes uniquely found by binary strategy; red: genes uniquely found by *DASE*; grey: genes found by both methods. **d)**. KEGG pathways (p < 0.05) enriched in the uniquely linked genes by *DASE*.

Beyond transcription factor binding, we examined the downstream effects of SE alterations on gene expression. We identified SE target genes in each cell type using 3D chromatin interactions based on POLR2A targeted ChIA-PET data. As expected, the gained CEs usually establish new gene targets, while the dismissed CEs remove existing targets (**Figure 3c**). Consequently, SEs with increased activity (e.g., strengthened or lengthened with gained CEs) in one cell type usually target more genes compared to their altered forms (e.g., weakened or shortened with dismissed CEs) in the other cell type (**Figure 3d**). Interestingly, we observed that such effects differed across the differential SE patterns, with heavier effects presented by *overall-change, shortened* and *hollowed* SEs, and nearly no effects by *shifted* SEs. The marginal effects by *shifted* SEs are expected as they provide no signs of the altering directions of SE activities. Here, to minimize the sequencing coverage effects on gene target counting with ChIA-PET data, we normalized the count differences by subtracting the median count difference (i.e., 1) of the control SE group (i.e., the non-differential SEs).

Differential analysis of gene expression between K562 and MCF7 cell lines indicated that the gained CEs between the two cell types were significantly associated with upregulated gene expression (**Figure 3e**). A similar association was also observed at the SE level, with increased SE activity presenting higher amplification on gene expression (**Figure 3f**). Again, *overall-change* and *shortened* SEs showed higher regulatory effects, while *shifted* SEs presented nearly no effects. Here, *hollowed* SEs showed no impact on gene expression, indicating their functions might be limited to maintaining the proper number of gene targets (**Figure 3d**). As a control, we observed no significant effects on gene expression by the non-differential SEs (**Figure 3f**).

We then performed pathway enrichment analysis on genes linked by different types of SE alterations. The majority of genes (∼75%) are linked by only one type of differential SEs (**Figure 3g**), with the *overall-change* and *shortened* SEs linked with the most and comparable number of genes. We focused on these genes linked by only one type of SEs and identified distinct sets of signaling pathways uniquely associated with each type of differential SEs (**Figure 3h**). For instance, FoxO Signaling Pathway (27-29), cAMP Signaling Pathway (30), and AMPK signaling pathway (31) are enriched with genes linked to *overall-change, shortened*, and *hollowed* SEs, respectively. In summary, different patterns of SE alterations synergistically build up gene regulation by playing distinct roles in modulating gene expression and cellular functions.

### Accounting for internal dynamics improves identification and interpretation of differential SEs

Besides the characterization of SE structural alterations, *DASE* presents an overall improvement on differential SE identification over the existing binary strategy (12-16). We summarized the improvements based on pair-wise comparisons across the six cancer cell types. These cells have H3K27Ac ChIP-seq, 3D chromatin interaction, and gene expression data available which enable the functional assessment on targeting gene expression. On average, the discrepant identification between *DASE* and the binary strategy accounts for ∼18% of the total SEs and covers all patterns of SE alterations (**Figure 4a**). Most newly identified differential SEs by *DASE* have structural alterations (∼91%). Among all discrepant differential SEs, the *overall-change* SEs newly identified by *DASE* showed the strongest impact on altering the numbers and expression of the gene targets (**Figure 4b**), suggesting they were falsely identified as non-differential by the binary strategy. The other newly identified SEs by *DASE* presented relatively higher gene effects compared to the differential SEs only identified by the binary strategy (**Figure 4b**), suggesting the overall improved sensitivity and specificity by the *DASE* identification. Here, we didn’t assess the *shifted* and *other complex scenario* SEs in this analysis as they provided no signs of the altering direction for gene expression. To avoid confounding effects from genes targeted by multiple SEs, we left out genes that were also linked by the common differential or non-differential SEs in the analysis.

We further evaluated *DASE* by gene functions linked to the differential SEs. We identified cell-type-specific regulated genes that were linked to the SEs with increased activity (i.e., overall increased, lengthened, or hollowed with increased CEs) in one cancer cell type compared to the other five cell types. We then compared the obtained gene list between *DASE* and the binary strategy. Surprisingly, *DASE* recovered nearly all the cell-type-specific regulated genes by the binary strategy and found additional genes mainly linked to the SE structural alterations (**Figure 4c**). We examined the pathways enriched in these additionally identified genes and found a number of cancer-type-specific pathways, indicating the cancer-specific roles of the novel structural alterations (**Figure 4d**). For example, Insulin Signaling Pathway (32), ErbB Signaling Pathway (33), Thyroid Hormone Signaling Pathway (34), TGF-beta Signaling Pathway (35), NF-kappa B signaling pathway (36), and Neurotrophin Signaling Pathway (37), linked to SEs that *DASE* uniquely identified in A549, HCT116, HepG2, K562, MCF7 and SK-N-SH, respectively. In summary, *DASE* showed improved sensitivity in linking differential SEs to cancer-specific regulation, particularly through the consideration of internal structural alterations.

### Accounting for internal dynamics maximize the discerning of cell identity

Given the improved sensitivity in the cross-cancer analysis above, we further evaluate *DASE* by within-cancer comparison. We applied *DASE* to compare SE profiles between two similar cancer cell lines, BC1 and BC3, that are B lymphocyte cells derived from Lymphoma under different viral infections. We previously demonstrated that different viral infections led to a distinct enhancer connectome on these cell lines (16).

Overall, the two similar cell lines presented much higher similarity of SE profiles (**Figure 5a**). We linked the differential SEs to their target genes using chromatin interactions identified by H3K27Ac HiChIP datasets. Similar gene effect patterns were observed across differential SE patterns, as we found previously (**Figure 3c-f**). The linked genes were enriched in both frequency and expression in the same direction as CEs/SEs altering between BC1 and BC3 cell lines (**Figure 5b-e**). Specifically, such effects are stronger by *overall-change* SEs, followed by *shortened* SEs, consistent with the findings in cross-cancer analysis (**Figure 3c-f**). Finally, we extracted the uniquely linked genes by the differential SE patterns (**Figure 5f**) and performed pathway enrichment analysis. We found unique pathways such as Viral Carcinogenesis particularly linked to the *shortened* SEs (**Figure 5g**). This suggests *shortened* SEs play key roles in gene dysregulation in response to the different viral carcinogenesis between BC1 and BC3 cell lines (38,39). Therefore, by accounting for the SE internal dynamics, we found that cell-line-specific gene regulation linked to differential SEs, particularly those with structural alterations, highlighting the differentiated roles of SE categories and the importance of featuring internal dynamics in SE differential analysis.

**Figure 5.**
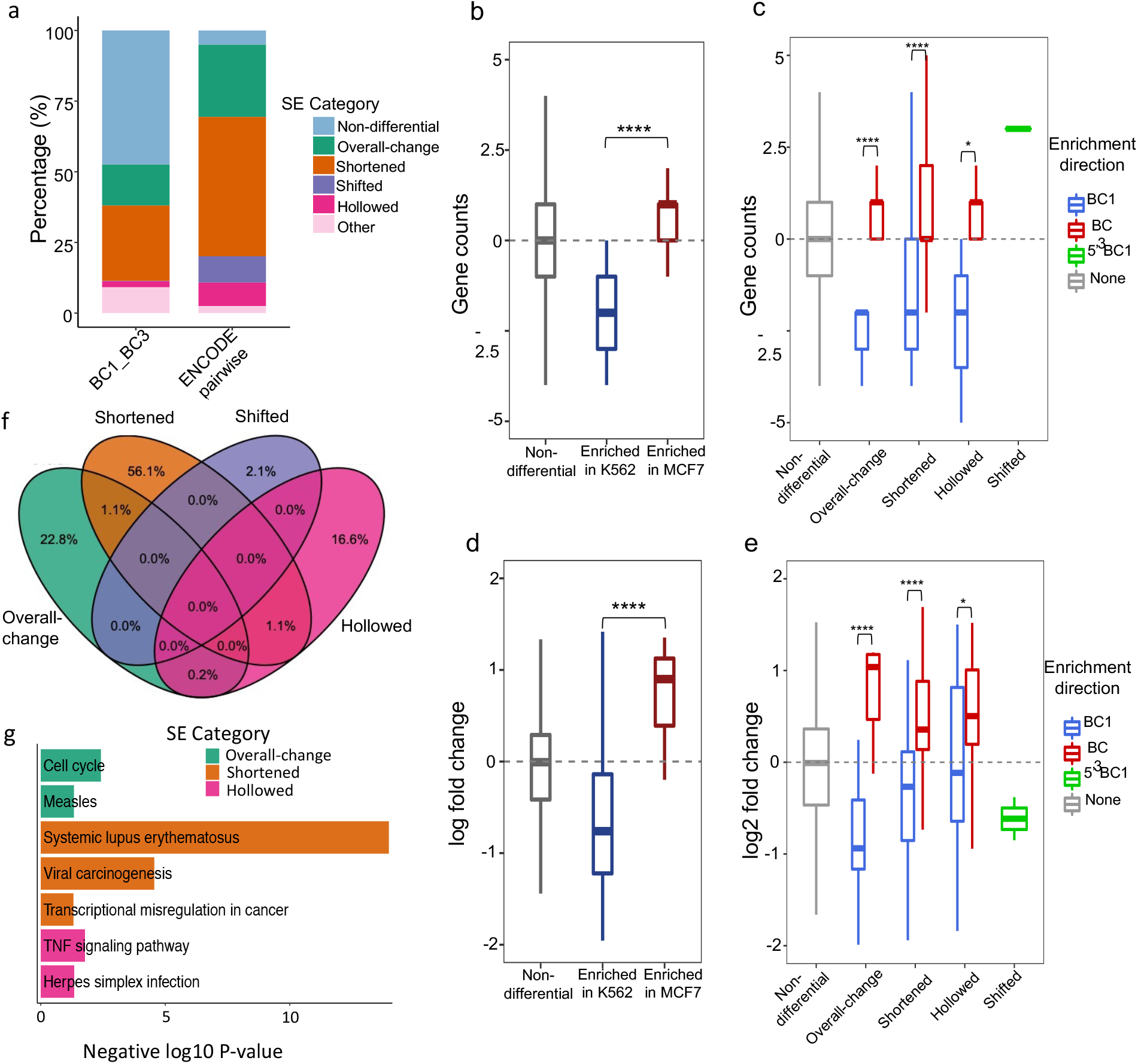
Identifying SE differences between similar cancer cell lines. **a)**. Proportions of differential SE categories identified from within-cancer (BC1 vs. BC3) and cross-cancer comparisons (pairwise between six cancer cell types from ENCODE). Count differences (**b**,**c**) and log2 fold-changes (**d**,**e**) of linked genes by differential CEs (**b**,**d**) and SEs (**c**,**e**) between BC1 and BC3 cell lines. Gene targets are identified based on chromatin contacts from H3K27Ac HiChIP data. **f)**. Overlaps of SE-linked genes across four differential SE categories between BC1 and BC3. **g**. KEGG pathways (p < 0.05) that uniquely associated to each differential SE category.

## DISCUSSION

In this manuscript, we proposed a novel computational method *DASE* to identify differential SEs by summarizing the internal dynamics. We categorized differential SEs into five major groups based on their overall activity and structural alterations: *overall-change, shortened, hollowed, shifted*, and *others*. By assessing differential SEs with the enriched transcription factors and linked target genes, we found distinct characteristics associated with different groups of SEs, such as linking with different numbers of genes and affecting gene expression at divergent impact. When compared with the widely adapted binary approach, *DASE* found an improved list of differential SEs which are linked to distinct cancer-specific gene functions. This highlighted the elevated performance by *DASE* identification. It further demonstrated the increased power in identifying cell-line-specific SE regulation when applied to similar cell lines.

Specifically, our improved performance is powered by the consideration of SE internal dynamics. For instance, SEs might show frequent internal alterations yet with no overall activity changes, as shown in our study. These differences, however, if not accounted for, could under-estimate the genome-wide variation of SE profiles and consequentially bias the evaluation of SE effects on gene regulation. On the other side, significant activity changes of SEs are usually combined with structural alterations, either globally or partially, indicating modeling structural differences won’t lose specificity in detecting true SE differences. However, we did notice that some SEs hold marginal activity changes which were weighted differently as discrepant calls between the binary strategy and our methods. Nevertheless, these SEs usually showed lower effects on gene expression compared to other differential SEs. Especially, those discrepant differential SEs could regulate genes in alternative way by altering the number of linked gene targets if they present significant structural alterations.

One limitation of our methods is that we cannot identify structural differences when a SE contains only one CE. We proposed a weighted spline model to account for the contribution of CEs by their width and activities. Thus, the model requires at least two CEs within a SE to generate a confident estimation. In practice, we identified SEs with only one CE as either non-differentials or *overall-change* if their activities are significantly altered. In addition, we identified differential SEs as *other complex scenarios* if their internal patterns cannot be attributed to all other categories. We detailed this in the ***Methods*** section. In practice, we found this category only accounts for a small portion of SEs (**Figure 5a**). We leave a closer interpretation of such complexity to future work. SEs were conceptionally defined based on the intensity and enrichment of consecutive enhancers (2,3). As a result, significant changes of SEs may correspond to two scenarios: activity changing between two SEs or status transitions between SEs and regular enhancers. These scenarios may associate with different functional interpretations since regular enhancers tend to regulate less and closer genes compared to SEs. Although we didn’t provide approaches to discriminate the two scenarios as that goes beyond the scope of our proposed study, feasible strategies could be implemented in future work to improve the downstream interpretation. For instance, scanning the distances between SEs and gene promoters could help filter regular enhancers as they are usually close to their gene targets (40,41). Also, SE alterations can be linked to the status changes of local chromatin, such as phase separation (42), to help determine if transitions occur between SE and regular enhancers. These require the integration of additional datasets to define chromatin statuses.

## MATERIALS AND METHODS

### Data acquisition

H3K27ac enrichment, gene expression and 3D interaction data were downloaded from ENCODE data portal (43) and GEO repositories (44). Specifically, quality-controlled alignment files of H3K27Ac ChIP-seq and RNA-seq, and chromatin contacts files of POLR2A ChIA-PET were downloaded from ENCODE for six selected cancer cell lines (A549 - Lung Cancer, HCT116 - Colorectal Cancer, HepG2 - Liver Cancer, K562 - Leukemia, MCF7 – Breast Cancer, and SK-N-SH - Neuroblastoma) (accession ID documented in **Supplementary Table 1**). Raw sequencing files of H3K27ac ChIP-seq, RNA-seq and H3K27ac HiChIP for BC1 and BC3 cell lines were downloaded from GEO with accession IDs GSE136090 (16) & GSE114791 (45) (**Supplementary Table 1**).

### H3K27Ac ChIP-seq data pre-processing

Raw ChIP-seq data from GEO was first aligned to human genome using Bowtie2 (46). Then, all alignment files were processed for peak calling using MACS2 (11), followed by SE detection using ROSE (2). All tools were applied with default parameters. ChIP-seq blacklist regions were excluded for downstream analysis (47).

### RNA-seq data analysis

RNA-seq alignment files downloaded from ENCODE were quantified for gene expression using featureCount (48) based on GENCODE annotations. Raw FASTQ files from GEO were processed with semi-alignment and quantification tool Salmon (49) to generate gene expression count table based on GENCODE transcriptome. Then, differential analysis of gene expression was estimated using DESeq2 (50) for all two-condition comparisons. The shrunk fold-changes were extracted to represent gene expression differences (51).

### 3D chromatin contacts analysis

The chromatin contacts generated by ENCODE project from POLR2A ChIA-PET data were directly adapted to link genes and super enhancers for ENCODE cancer cell lines. Basically, ENCODE project applied strict quality controls, and filtered confident chromatin contacts with at least 3 normalized interactions (52). H3K27Ac HiChIP data of BC1 and BC3 cell lines are analyzed the same as previously described (16). In brief, reads were aligned to human genome using HiC-Pro (53). Sequencing replicates were merged to call chromatin contacts using hichipper (54) with confident interactions defined as at least 3 normalized interactions.

### Differential analysis of CEs and binary SE differences

For each comparison between two cell lines that both have two ChIP-seq replicates, a uniform peak list was first created by merging overlapped peaks across the compared samples. ChIP-seq reads were then quantified using featureCount (48) to generate a read count table for the peak list. Differential peak analysis was performed by adapting DESeq2 (50) (with parameter *type=‘mean’*) to account for the varied dispersion between peaks with low and high read counts. The differential statuses of CEs (H3K27Ac peaks within SEs) were extracted based on their estimated log2 fold-changes and corresponding q-values. We also extracted the normalized coverage for CEs as the weight inputs for SE differential analysis.

Binary SE differences were estimated based on the presence and absence of SEs between compared conditions. Basically, if a SE presents in both compared conditions at the same given location regardless size or activity changes, it will be identified as non-differential. In contrast, if a SE only presents in one condition at a given location, it will be identified as differential.

### Modeling differential SEs with SE internal dynamics by DASE

*DASE* identifies differential SEs by accounting for the combinatorial effects of CEs weighted with their activities and locations. In detail, the methods include the following steps (**Supplementary Figure 5**).

#### Input preparation

A uniform list of SEs is generated by merging overlapped SEs between compared conditions. The differential statuses (log2 fold-change) of all CEs located within SEs are extracted as well as their activities (ChIP-seq coverage) and locations (genomic coordinates), as calculated above. In practice, we select the maximum ChIP-seq coverage between compared conditions for each CE to provide better weights in the spline model below.

#### Weighted spline model

For each SE, the log2 fold-change values of CEs stratified by their genomic locations are fitted using b-spline model, where the importance of CEs is weighted by their relative activities. As a result, CEs show less impacts on the spline fitting if they have low activities and stay close to other CEs. We implement the spline model using R package *splines*. In addition, to ensure the robustness of b-splines in the case of too many low- or mild-activity CEs, we pre-estimate the degree of freedom for each fitting based on the number of top ranked CEs in each SEs. We choose top ranked CEs as the minimum number of enhancers that build up at least 95% of total SE activity. In detail, we set the degree of freedom of b-spline as 2, 3 and 4 if this number of top ranked CEs is less than 4, between 4-6, and larger than 6, respectively.

#### Significance estimation

We use permutations to define significant fitted values by b-spline. In brief, we randomly shuffle enhance activities in each compared sample, re-estimate the differential statuses of all CEs and re-fit splines for all SEs. As a result, we generate a null distribution of fitted b-spline values for all CEs. We repeat the processes 10 times for a stable null distribution. Significant fitted values are defined as those having greater or smaller values than the upper or lower inflection points (significant thresholds) of the null distribution (**Supplementary Figure 5**).

#### Status summarization for SE sub-regions

We divide each SE into multiple sub-regions using the intersects of b-spline curves and the significant thresholds (**Figure 2**). For instance, the curve located above the upper threshold indicates an up-altered partial region within the SE, while the curve located below the lower threshold indicates a down-altered partial region. The curves in between indicate non-altered SE sub-regions. To decrease potential noises in SE segmentation, we ignore sub-regions in which CEs account for less than 1% of the SE activities.

#### Overall differential status

We further summarize the overall differential statuses for SEs with heuristic approaches based on the statuses, locations, and activity occupancies (*i*.*e*., the percentage of activity over the total activity of SEs) of segmented sub-regions. Specifically, if only one region is resulted from segmentation of b-spline curve of a SE, the SE will be identified as either non-differential or differential depending on the status of that segment. If two segments are resulted (i.*e* one is significant and one is un-altered), we determine differential SEs based on the activity occupancy of the significant segment. In detail, two-segment SEs are identified as non-differential, *shortened* or *overall-change* if the significant segment occupies less than 10%, between 10%-90% and more than 90% of total SE activities. For a three-segment SE, we first check if it is *hollowed* based on whether the middle segment is significantly altered. If not, we check if it is *shifted* based on whether the three segments cover three different statues (i.e., up-altered, down-altered and on-altered) separately. Otherwise, the remaining three-segment SEs fall into the following situation: the middle segment is non-significant while the left and right segments are both significant with the same altering direction. We then identify the overall SE statues as non-differential, *shortened* and *overall-change* based on the total activity occupancies of the left and right segments as below 10%, between 10%-50% and above 50%. It is noted that the *overall-change* are filtered with different criteria (break points at 90% vs 50%) in two-segment and three-segment SEs, to account for the total size impacts from the altered segments. For a SE with more than 3 segments, it is identified as *other complex scenario* except that a four-segment SE holding all three statuses is defined as *hollowed*. Finally, we rank the significance of differential SEs using the activity occupancies of the significant segments separately for *overall-change, shortened, hollowed* and *shifted* SEs.

### Transcription factor enrichment analysis

ChIP-seq bam files for 78 documented transcription factors in both K562 and MCF7 cell lines were downloaded from ENCODE with accession ID provided in **Supplementary Table 1**. After calling differential SEs between MCF7 and K562 with *DASE*, we extracted transcription factor occupancy from ChIP-seq data for significant differential (fold-change >4 & q-value < 0.05) CEs that locate within differential SE categories: *overall-change, shortened, hollowed*, and *shifted*. The occupancy heatmap for transcription factors were generated with Deeptools v3.5.1 (55).

### SE-gene targeting

We identify SE-gene targeting relationship using 3D chromatin contacts generated from POLR2A ChIA-PET or H3K27Ac HiChIP. Basically, a valid targeting is defined if one end of chromatin contacts is overlapped with SEs, while the other end is overlapped with gene promoters (selected as -3kbp - 1kbp from genes’ transcription start sites). Targeting relations are ignored if the SE-promoter distances are less than 20kb or greater than 500kb.

### Pathway enrichment analysis

For pathway enrichment in genes linked by different SE categories, gene sets were first identified for each SE categories based on SE-gene targeting relations in both compared conditions. Then, only uniquely linked genes by each SE category were selected for pathway enrichment using DAVID Bioinformatics Resources v6.8 (56) based on KEGG database (57). For pathway enrichment in genes linked by cancer-specific SEs, genes were selected as those only identified by *DASE* compared to the binary strategy. Significant pathways were selected to have p-value less than 0.05.

