## Supplemental Figures for "Summarizing internal dynamics boosts differential analysis and functional interpretation of super enhancers"

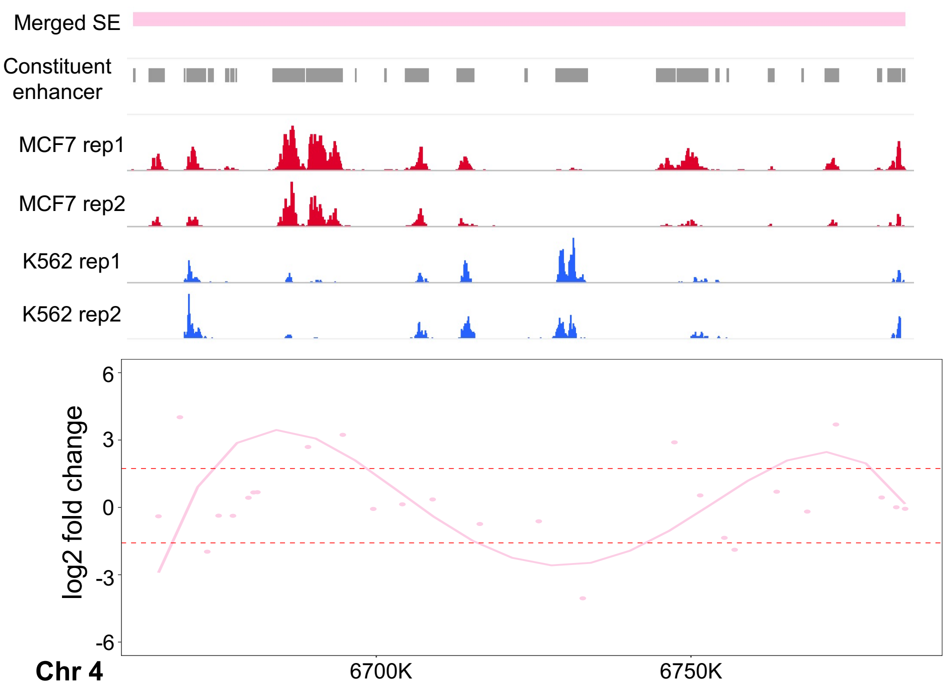


**Supplementary Figure 1. An example of *Other complex* scenario SEs.** The upper panel lists in order are the SE regions, CEs, H3K27Ac ChIP-seq coverage in two cell types with two replicates. The lower panel shows the fitted b-splines in addition to the original log2 fold-change values for CEs (points). Dashed lines indicate the estimated thresholds from permutation to define differential segments within SE regions.

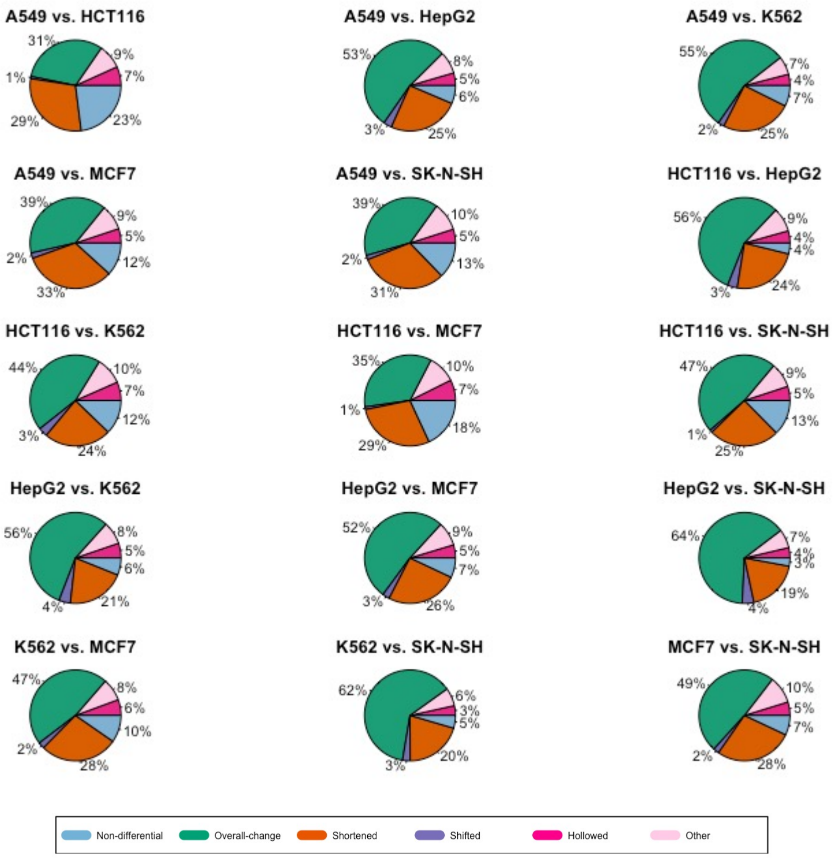


**Supplementary Figure 2. Percentages of the estimated differential SE categories in pair-wise comparisons of the six cancer cell types.** Compared cell types are labeled as the title in each sub-figure.

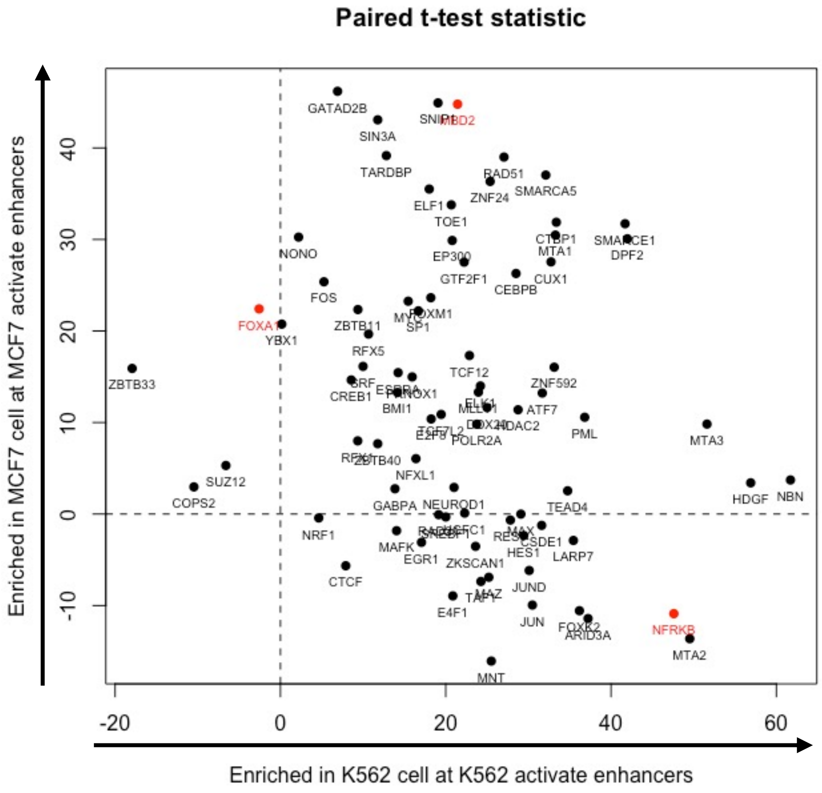


**Supplementary Figure 3. Comparisons of transcription factor enrichment at the differential CEs between K562 and MCF7.** X-axis indicates the paired t-test statistics for transcription factor binding signals between K562 and MCF7 at the K562 activate enhancers. Y-axis indicates the paired t-test statistics for transcription factor binding signals between K562 and MCF7 at the MCF7 activate enhancers. Transcription factor binding signals were extracted as the H3K27Ac ChIP-seq signals at the middle base pair of the enhancers defined by ChIP-seq peaks. Each points represents a transcription factor. Red points (FOXA1, NFRKB, and MBD2) are the selected examples in the main manuscript.

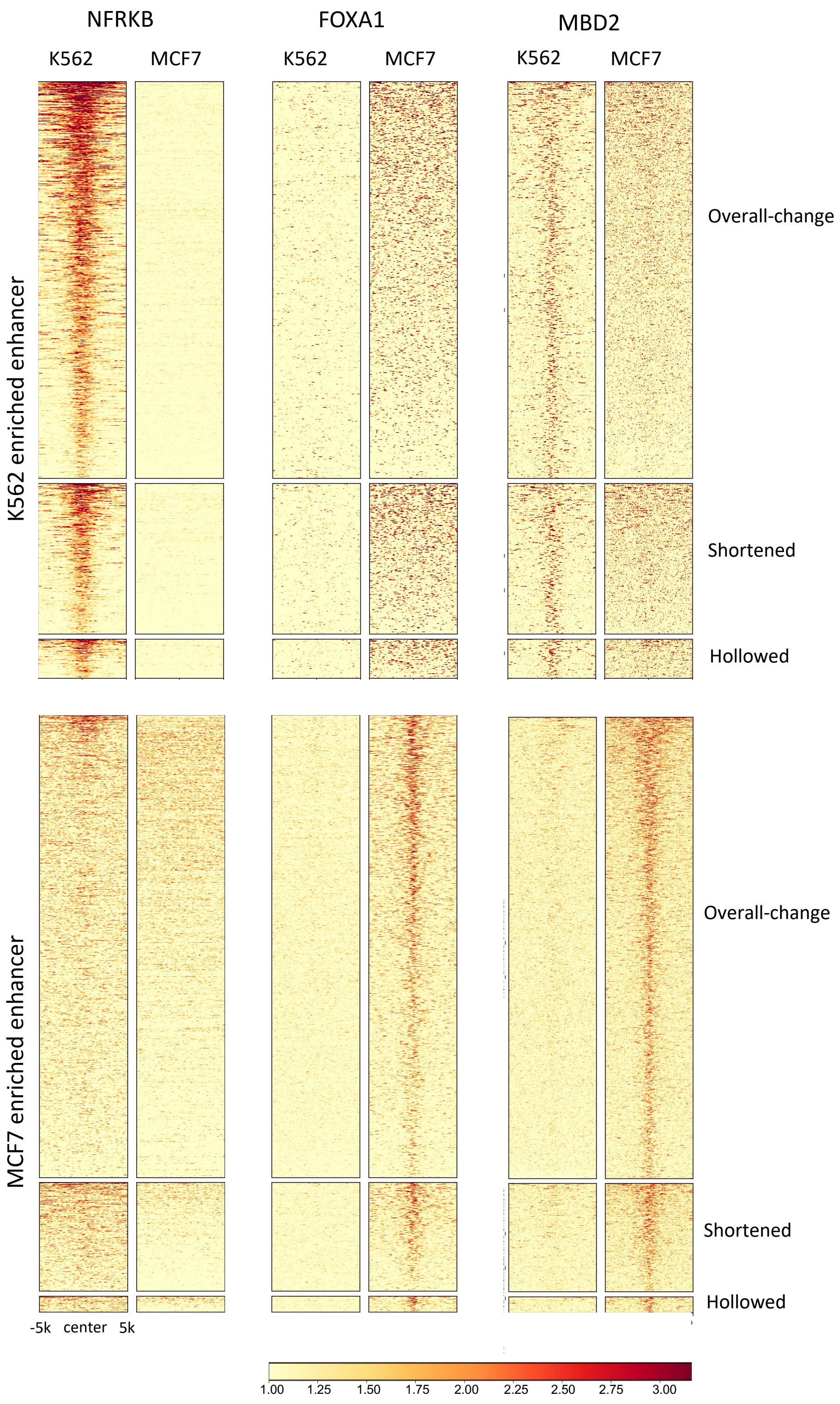


**Supplementary Figure 4. Transcription factor enrichment at the differential CEs from three categories of differential SEs.** The heatmaps are plotted the same as Figure 3b except signals at constituent enhancers of different SE categories are separated.

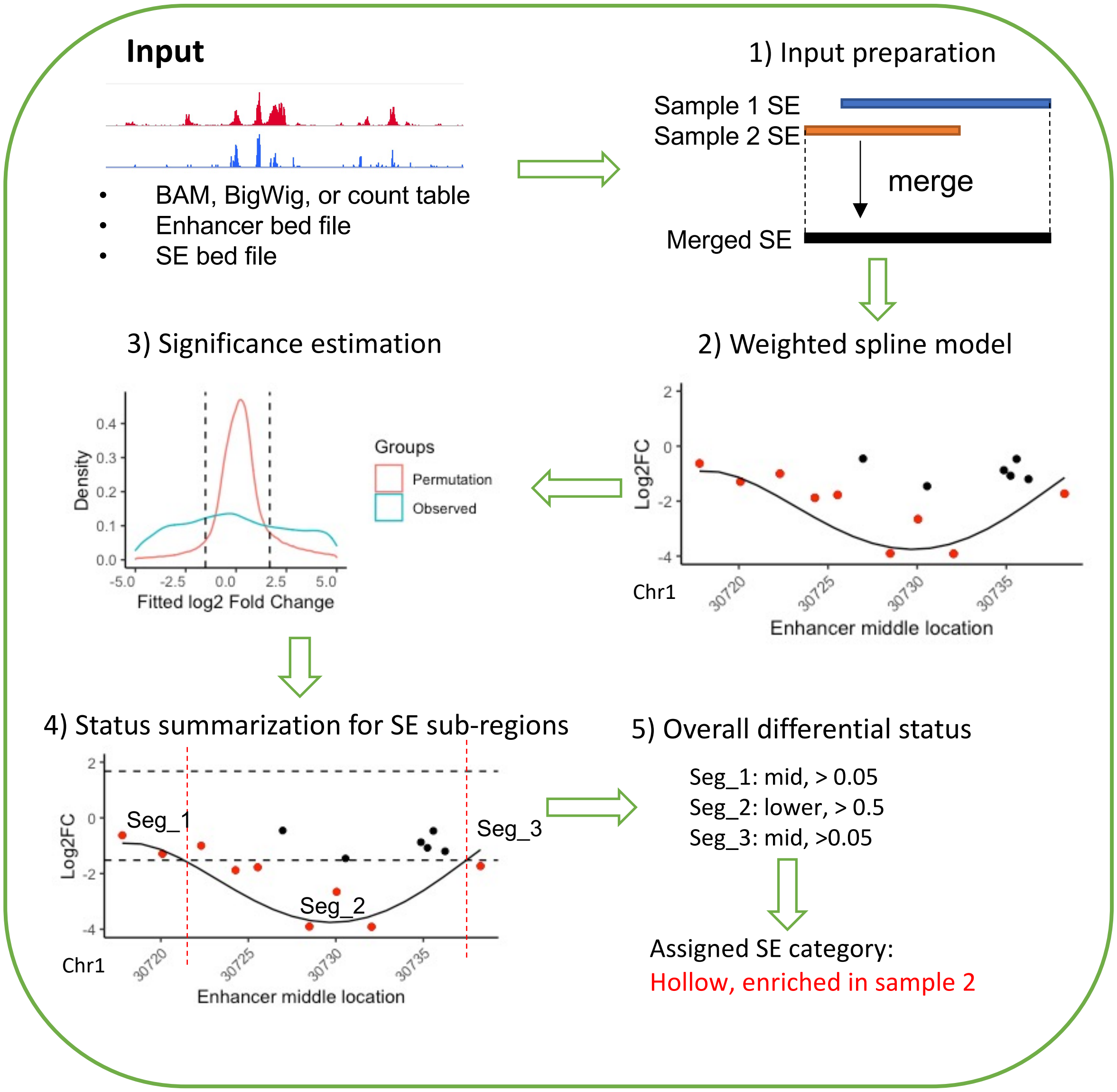


**Supplementary Figure 5. Workflow of *DASE*. Input** shows the required input files by *DASE*, including enhancer activity file (ChIP-seq BAM, BigWig, or count table), enhancer location file (BED), and SE location file (BED). Five main processing steps by *DASE* are demonstrates in the following order, including input preparation, weighted spline model, significance estimation, status summarization for SE sub-regions, and overall differential status.
